# Advantages of Multi-shell Diffusion for Studies of Brain Development in Youth

**DOI:** 10.1101/611590

**Authors:** Adam R. Pines, Matthew Cieslak, Graham L. Baum, Philip A. Cook, Azeez Adebimpe, Diego G. Dávila, Mark A. Elliott, Robert Jirsaraie, Kristin Murtha, Desmond J. Oathes, Kayla Piiwaa, Adon F. G. Rosen, Sage Rush, Russell T. Shinohara, Danielle S. Bassett, David R. Roalf, Theodore D. Satterthwaite

## Abstract

Diffusion tensor imaging (DTI) has advanced our understanding of how brain microstructure evolves over development. However, the proliferation of multi-shell diffusion imaging sequences has coincided with notable advances in the modeling of neuronal diffusion patterns, such as Neurite Orientation Dispersion and Density Imaging (NODDI) and Laplacian-regularized Mean Apparent Propagator MRI (MAPL). The relative utility of these newer diffusion models for understanding brain maturation remains sparsely investigated. Additionally, despite evidence that motion artifact is a major confound for studies of development, the relative vulnerability of these models to in-scanner motion has not been described. Accordingly, in a sample of 123 youth (ages 12-30) we evaluated DTI, NODDI, and MAPL for associations with age and in-scanner head motion at multiple scales, including mean white matter values, voxelwise analyses, and tractography-based structural brain networks. Our results reveal that multi-shell diffusion imaging sequences can be leveraged to robustly characterize neurodevelopment, even within the framework of DTI. However, these metrics of diffusion are variably impacted by motion, highlighting the importance of modeling choices for studies of movement-prone populations. Our findings suggest that while traditional DTI is sensitive to neurodevelopmental trends, contemporary modeling techniques confer key advantages for neurodevelopmental inquiries.

## 1. INTRODUCTION

Diffusion-weighted imaging (DWI) has informed our understanding of both local tissue (Basser and Pierpaoli 1996; Koh and Padhani 2006; Svolos et al., 2014) and distributed network properties of the brain *in vivo* (Sporns, Tononi, and Kötter 2005; Gollo et al., 2018). DWI has proven to be particularly useful for studying neurodevelopment, and has provided critical evidence of the protracted maturation of white matter from infancy into adulthood (Lebel et al., 2008; Schmithorst and Yuan 2010). Recent studies have leveraged tools from network neuroscience and established that structural networks reconfigure in development to promote efficient communication (Hagmann et al., 2010; Fan et al., 2011; Grayson et al., 2014; Baum et al., 2017; Uddin et al., 2011; Baker et al., 2015; Bassett, Zurn, & Gold, 2018; Huang et al., 2015).

Most DWI studies have used diffusion tensor imaging (DTI) with a single diffusion weighting to characterize observed diffusion patterns as indices of neuronal microstructure (Lebel & Deoni, 2018; Lebel, Treit, & Beaulieu, 2017). While valuable, these studies may have been limited by certain characteristics of diffusion tensor model and single-“shell” imaging. In practice, DTI-derived metrics underestimate diffusion restriction in voxels within crossing fibers (Jeurissen et al., 2013; Jones and Cercignani 2010; Volz, Cieslak, and Grafton 2018; De Santis et al., 2014) and are systematically impacted by in-scanner motion, which is often prominent in children (Yendiki et al., 2014; Ling et al., 2012; Baum et al., 2018; Roalf et al., 2016). More recently, a new generation of models have been developed to leverage multiple *b*-values (“shells”). When systematically varied over a DWI acquisition, the differential tissue responses elicited by different *b*-values can be used to model more detailed features of the cellular environment (Stanisz et al., 1997; Clark, Hedehus, and Moseley 2002; Assaf and Basser 2005). These models can be broadly separated into “tissue” and “signal” models (Alexander et al., 2017; Ferizi et al., 2017): tissue models attempt to classify signal attributable to different classes of biological tissues, while signal models do not attempt to delineate tissue classes.

Although several tissue models were foundational for multi-compartment modeling of diffusion images (Assaf and Basser 2005; Alexander et al., 2010), Neurite Orientation Dispersion and Density Imaging has become the most widely used (NODDI; Zhang et al., 2012). NODDI provides estimates of the directional distribution of neurites (axons and dendrites), as well as compartmental volume fractions. These include the proportion of volume estimated to be intracellular, extracellular and isotropic in each voxel based on the estimated contributions of these compartments to the diffusion signal. Recent work suggests that NODDI may be more attuned to biological features of brain development than DTI (Chang et al., 2015; Genc et al., 2017; Nazeri et al., 2015; Mah, Geeraert, and Lebel 2017; Eaton-Rosen et al., 2015; Deligianni et al., 2016; Ota et al., 2017). However, it remains unclear as to how useful NODDI-based measures are for studies of brain networks, or how they are impacted by in-scanner motion.

In contrast to tissue-based models like NODDI, “signal” based methods use the Fourier relationship between *q*-space signal and the ensemble average propagator (EAP) to characterize the intra-voxel diffusion process. Two recently-introduced techniques which model the EAP are Mean Apparent Propagator MRI (MAP-MRI; Özarslan et al., 2013) and Laplacian-regularized MAP-MRI (MAPL; Fick et al., 2016). These models allow estimation of the likelihood of water molecules undergoing zero net displacement in in up to three dimensions in any voxel (Özarslan et al., 2013). In contrast to the accumulating number of studies which have used NODDI to study brain development, MAPL has not been previously used in studies of brain maturation. Furthermore, like NODDI, it remains unknown how motion may impact EAP-based measures.

Here we sought to describe the relationship between three diffusion models and both brain development and in-scanner motion. We evaluated how diffusion metrics from DTI, NODDI, and MAPL are associated with both age and in-scanner motion in a sample of 123 youth who completed multi-shell diffusion imaging. Importantly, we included DTI metrics derived from solely the *b*=800 shell (to more closely match a traditional DTI scan), as well as the full multi-shell scheme. Statistical associations were examined across multiple scales of analysis, including mean white matter values, voxelwise analyses, and tractography-based networks. As described below, we present new evidence that multi-shell diffusion acquisitions can be leveraged to provide advantages for studies of the developing brain.

## 2. METHODS

### 2.1 Participant characteristics

After quality assurance exclusions (section 2.3), we studied 123 participants between the ages of 12 and 30 years old(*M* = 21.08, *SD* = 3.54, 70 females). Potential participants were also excluded for metallic implants, claustrophobia, pregnancy, acute intoxication, as well as significant medical and/or developmental conditions that would have impeded participation. All subjects were compensated for their time, and all protocols were approved by the University of Pennsylvania’s Institutional Review Board.

### 2.2 Image acquisition

All participants were imaged on a 3-Tesla Siemens MAGNETOM Prisma with a T1-weighted structural and diffusion-weighted scan. Our structural scan was a 3 min 28 s MPRAGE sequence with 0.9 x 0.9 x 1.0 mm^3^ resolution (TR = 1810 ms, TE = 3.45 ms, inversion time = 1100 ms, flip angle = 9 degrees, acceleration factor = 2). Our DWI sequence was a single-shot, multiband, multi-shell acquisition protocol (TR = 3027 ms, TE = 82.80 ms, flip angle = 78 degrees, voxel size = 1.5 mm^3^ isotropic, FOV = 210 mm, acquisition time = 6 minutes 12 seconds, acceleration factor = 4, phase-encoding direction = anterior to posterior) with 3 diffusion-weighted shells at b=300 s/mm^2^ (15 volumes), b=800 s/mm^2^ (30 volumes), and b = 2000 s/mm^2^ (64 volumes). The sequence included 9 b=0 s/mm^2^ scans interspersed throughout. We also acquired a b=0 s/mm^2^ reference scan with the opposite phase-encoding direction (posterior to anterior) to correct for phase-encoding direction-induced distortions.

### 2.3 Pre-processing and quality assurance

Distortions induced by phase encoding were corrected using *topup* from the FMRIB Software Library (*FSL*; Jenkinson et al., 2012). Eddy-current distortions and in-scanner movement were corrected using *eddy* from FSL version 5.0.11 with both single slice and multiband outlier replacement (Jenkinson et al., 2012; Andersson et al., 2016; Andersson et al., 2017); this processing step also rotated the initial *b*-vectors from our sequence to align with estimated subject head motions. Following prior work, we quantified in-scanner motion using the root mean squared displacement over the course of the scan (mean relative RMS; Baum et al., 2018; Roalf et al., 2016). Mean relative RMS displacement was calculated between uncorrected interspersed b=0 images using publicly available scripts (https://www.med.upenn.edu/cmroi/qascripts.html). Following manual inspection of all T1 images, one subject was removed for poor T1 image quality. Additionally, one subject was removed for poor DWI quality (motion + 6.92 SD above the mean). Motion, distortion, and eddy-corrected images served as the common input to all diffusion modeling techniques.

### 2.4 Overview of diffusion metrics

Our evaluations include 14 diffusion metrics from three DWI modeling techniques. From DTI, we calculated fractional anisotropy (FA), mean diffusivity (MD), axial diffusivity (AD) and radial diffusivity (RD; Basser, Mattiello, and Le Bihan 1994). In accordance with previous applications of DTI to multi-shell data, we fit the DTI model using only the shell we expected gaussian diffusion responses (*b* = 800). Importantly, we also fit a DTI model to the entire multi-shell dataset using an iteratively reweighted linear least squares estimator tensor fit, (Veraart et al., 2013) which yielded an equivalent 4 diffusion metrics of interest. From NODDI, we calculated orientation dispersion indices (ODI), as well as intracellular (ICVF), and isotropic volume fraction (ISOVF) (Zhang et al., 2012). From MAPL, we evaluated return-to-origin (RTOP), return-to-axis (RTAP), and return-to-plane (RTPP) probabilities (Özarslan et al., 2013, Fick et al., 2016).

### 2.4.1 DTI metrics

DTI assesses the directionality and magnitude of water diffusion by assuming a Gaussian diffusion process in each voxel. DTI utilizes a 6 degrees of freedom symmetric tensor model fit to the observed signal. Subsequently, the primary direction of diffusion in a voxel is calculated by finding the largest eigenvalue of the tensor. After tensors are fit to a voxel, FA, MD, AD, and RD can be calculated from the corresponding eigenvalues. While MD is the averaged sum of these eigenvalues (representing the average magnitude of water diffusion), AD is derived from only the largest eigenvalue (representing the primary direction of diffusion). RD is the average of the remaining two eigenvalues, both representing eigenvectors orthogonal to the primary one. Finally, FA evaluates the magnitude of the eigenvalue associated with the primary direction of diffusion *relative* to the remaining eigenvalues. FA represents the fraction of anisotropy in a voxel aligned with a primary direction of diffusion. As diffusion shows increasing directional preference, FA increases (Soares, Marques, Alves, & Sousa, 2013; Basser, Mattiello, and Le Bihan 1994).

All DTI metrics were calculated in MRtrix3 using an iteratively reweighted linear tensor fitting procedure (Tournier et al., 2012; Veraart et al., 2013). As mentioned, we included FA, MD, AD, and RD derived from a DTI fit to all of the shells, as well as the same DTI metrics derived from the *b* = 800 shell only. This processing choice was made to account for the possibility that the utility of including more diffusion directions was outweighed by the non-Gaussian contribution of high *b*-value acquisitions.

### 2.4.2 NODDI metrics

NODDI estimates the directional distribution of neurites (axons and dendrites) in a voxel, and then matches diffusion patterns to that distribution. Like DTI, this model is informed by hindrance of diffusion unaligned with neuronal fibers, and unhindered diffusion along their prominent axes. Unlike DTI, the introduction of a 3D neurite distribution allows for modeling diffusion restriction in fiber populations with dispersed orientations.

NODDI attempts to parse the diffusion signal into discrete contributions of cellular compartments. The total signal is set to equal the sum of the contributions from each compartment, such that A = (1 − *V*_*iso*_)(*V*_*ic*_*A*_*ic*_ + (1 − *V*_*ic*_)*A*_*ec*_)+ *V*_*iso*_*A*_*iso*_, where *A* is the full diffusion signal, *A*_*ic*_, *A*_*ec*_, and *A*_*iso*_ are the signal attributable to the intracellular, extracellular, and isotropic compartments, and *V*_*iso*_, *V*_*ic*_, and *V*_*ec*_ represent the fraction of tissue volume attributable to the corresponding compartments. In order to assign diffusion signal to one of these compartments, the method assumes neurites can be modeled as zero-radius cylinders, or sticks. NODDI then fits an estimated distribution of these sticks to a spherical distribution, which captures the estimated spread of neurite orientations. ODI measures this spread, which ranges from 0 (non-dispersed) to 1 (highly dispersed). A_ic_ is calculated with respect to this posited orientation dispersion in any given voxel. Intracellular signal is estimated by comparing the spherical distribution of neurite orientations with the distribution of unimpeded diffusion, yielding *V*_*ic*_, or the ICVF metric. Isotropic diffusion signal is attributed to a cerebrospinal fluid compartment, which yields the ISOVF metric (Zhang et al., 2012). Recent advances have markedly accelerated fitting the NODDI model; here we calculated NODDI using AMICO, which has been shown to accelerate fitting the NODDI model by several orders of magnitude without substantially impacting accuracy (Daducci et al., 2015).

### 2.4.3 MAPL metrics

Unlike tissue-based models such as NODDI, signal-based techniques seek to model the EAP directly and do not assume the separability of specific tissue compartments. Notably, the EAP is not limited to representing ellipsoids, and can therefore in theory capture arbitrary fiber configurations. MAP-MRI expresses MR signal utilizing Hermite functions as a basis set, which allows for rapid convergence of function solutions (Özarslan et al., 2013, Walter 1977). Building on MAP-MRI, Fick et al., (2016) recently introduced Laplacian-regularized MAP-MRI (MAPL). MAPL imposes additional smoothness on MAP-MRI’s coefficient estimation using the norm of the Laplacian of the reconstructed signal. This approach effectively penalizes model fits with physiologically improbable high local variances, which are more likely to be artifactual than reflective of signals of interest (Descoteaux et al., 2007). The authors also demonstrated that this method reduces error over MAP-MRI in voxels with crossing fibers (Fick et al., 2016).

MAP-MRI and MAPL allow for quantification of the likelihood that diffusing molecules undergo zero net displacement in one, two, or three dimensions. More specifically, RTOP estimates the probability of molecules undergoing no net displacement in any direction. Similarly, RTAP estimates the probability that molecules undergo no net displacement from their primary axis of diffusion, while allowing for displacement along that axis. Finally, RTPP estimates the probability that molecules are not displaced from their original plane perpendicular to the primary direction of diffusion, while allowing for movement of molecules within that plane (Özarslan et al., 2013). We fit the MAPL model with a radial order of 8, without anisotropic scaling, using generalized cross-validation for determining optimal regularization weighting. We conducted model fitting and generated RTOP, RTAP, and RTPP with dipy, an open-source diffusion imaging toolbox in Python (Fick et al., 2016; Garyfallidis et al., 2014).

### 2.6 Structural Image Processing

T1 images were processed using the ANTs Cortical Thickness Pipeline (Tustison et al., 2014). Images were bias field corrected using N4ITK (Tustison et al., 2010), and brains were extracted from T1 images using study-specific tissue priors (Avants, Tustison, Wu, et al., 2011). We utilized a custom young-adult template constructed via the *buildtemplateparallel* procedure in ANTs (Avants, Tustison, Song, et al., 2011). A custom template was used due to evidence demonstrating the utility of custom templates in reducing registration biases (Tustison et al., 2014). The T1 to template affine and diffeomorphic transformations were calculated with the top-performing symmetric diffeomorphic normalization (SyN) tool in ANTs (Klein et al., 2009). The transforms between T1 and the initial *b*=0 DWI images were calculated using boundary-based registration with 6 degrees of freedom (Greve and Fischl 2009). All transforms were concatenated so that only one interpolation was performed.

### 2.7 Network construction

Accumulating evidence suggests that structural brain networks undergo substantial maturation during youth (Hagmann et al., 2010; Fan et al., 2011; Grayson et al., 2014; Baum et al., 2017; Uddin et al., 2011; Baker et al., 2015). Accordingly, in addition to analysis of summary measures and scalar maps, we evaluated each measure in the context of structural networks. Subject brains were divided into network nodes using the Schaefer 200 cortical parcellation (Schaefer et al., 2014). The parcellation was warped to the custom template, and then projected back to each subject’s T1 and native diffusion space using the inverse of each transform. Whole-brain connectomes were constructed by representing each of the 200 regions as a network node, and by representing deterministic tractography streamlines as network edges. Tractography was conducted in Camino (Cook et al., 2006) using the Euler tracking algorithm in native diffusion space (Basser et al., 2000). The intersection between gray and 1mm-dilated white matter was used as both seed regions and termination points for tractography. We used voxels defined as CSF by the segmented T1 image as termination boundaries for streamlines. Voxels defined as white matter by the segmented T1 image were used as an inclusion mask for streamlines, ensuring that streamlines had to pass through white matter. Additionally, we imposed a curvature restriction on all streamlines. Fibers determined to curve more than 60 degrees over a 5 millimeter interval were discarded in order to mitigate the impact of noise on tractography (Bastiani et al., 2012). Lastly, the mean value of each diffusion metric was calculated along each edge in this network; these values were used as edge weights between nodes connected via tractography streamlines. However, ODI and ISOVF both represent deviation from diffusion-informed structural uniformity, maps consisting of 1 minus ODI and 1 minus ISOVF were utilized for weighted structural networks. Similarly, as higher MD and RD are both indicative of less uniform local microstructure, maps consisting of their inverse (1/RD and 1/MD) were utilized to evaluate their utility for weighting structural networks.

### 2.8 Statistical analyses

Within subjects, Spearman’s ρ between each diffusion metric for all white matter voxels was calculated. This involved masking diffusion images, vectorizing scalar values for each voxel within that mask, and correlating the vectors for each image. Subsequent analyses utilized consistent models across three levels of features (see schematic in **Figure 1**). First, we compared mean values within a white matter mask. Second, we conducted mass-univariate voxelwise analyses within white matter. Third, we evaluated tractography-based structural networks. At each level, in order to rigorously model linear and nonlinear associations with age, we used generalized additive models (GAMs; Wood, 2001, 2004) with penalized splines in R (Version 3.5.1) using the *mgcv* package (R Development Core Team, 2010; Wood, 2011). To avoid over-fitting, nonlinearity was penalized using restricted maximum likelihood (REML). Age was modeled as a penalized spline; sex and in-scanner motion were included as linear effects. To estimate in-scanner motion relations, we regressed out the effects of age and sex estimated from these models and evaluated the correlation between model residuals and head motion. This allowed us to quantify the effect size of the relationships between diffusion metrics and head motion of while controlling for other common sources of variance. Type I error was accounted for using the False Discovery Rate (FDR; Q<0.05).

**Figure 1:**
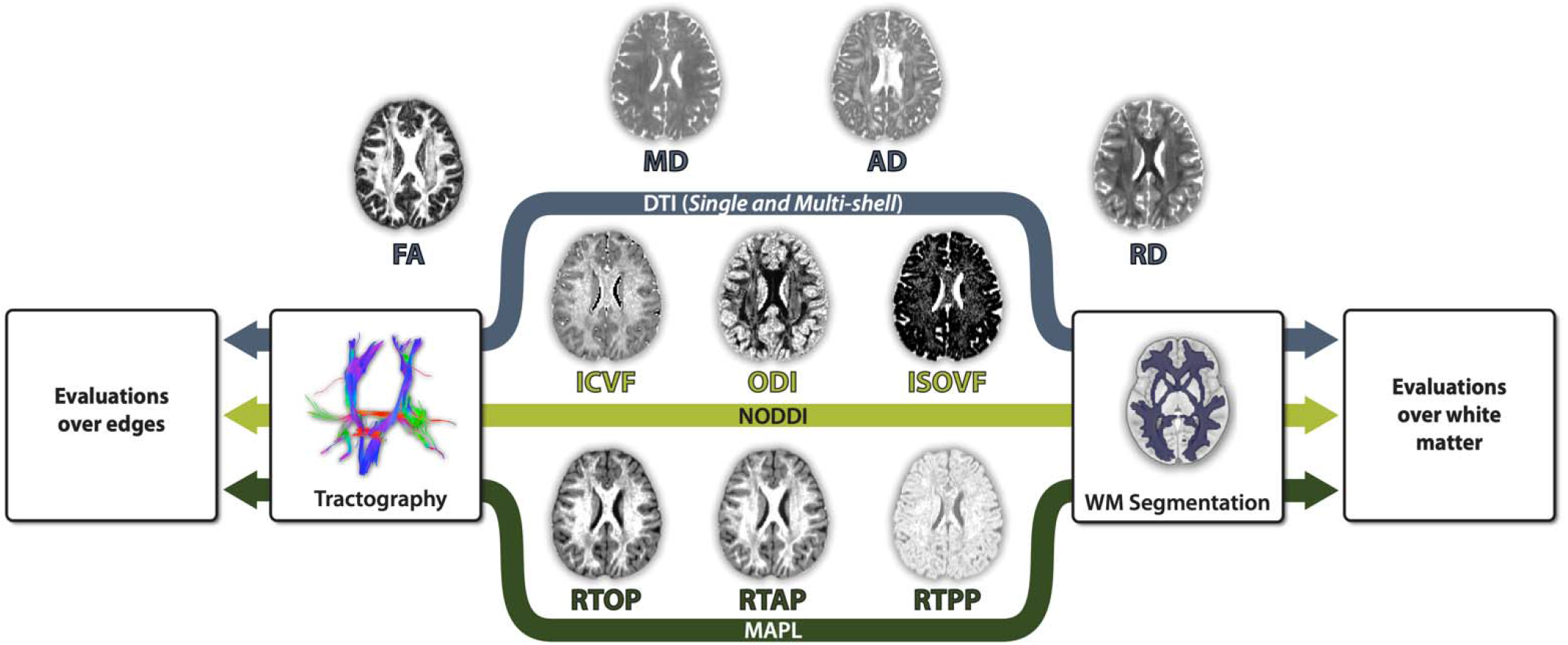
Analytic workflow. The DTI, NODDI, and MAPL models were fit to the same motion, distortion, and eddy-current corrected images, with the exception of the single-shell DTI fit, which only utilized our corrected *b* = 800 acquisitions. The resulting scalar maps were evaluated for associations with both age and data quality at multiple levels of analysis, including mean white matter values, mass-univariate voxelwise analyses, and analyses of network edges reconstructed by deterministic tractography (and then weighted by each measure).

### 2.9 Code availability

All analysis code used is available at: https://github.com/PennBBL/multishell_diffusion

## 3. RESULTS

### 3.1 Measures of diffusion show differential patterns of covariance

As an initial step, we investigated the relationships between all diffusion metrics of interest with Spearman’s correlations within white matter, and averaged those correlations across participants. This included correlations obtained when using a multi-shell DTI fit (**Figure 2A**, top triangle), and single-shell DTI fits (**Figure 2A**, bottom triangle). Some metrics of diffusion restriction were highly correlated with each other (i.e., FA, ICVF, and RTOP), and negatively correlated with metrics of diffusion dispersion (i.e., MD, ODI). Other metrics, like RTPP, demonstrated less systemic covariation with other metrics of interest, indicating that they may capture unique microstructural information. Multi- and single-shell DTI metrics also demonstrated high correlations to each other (Figure 2B), with MD being the least similar (*r* = 0.73). Next, we sought to understand the differential utility of these measures of diffusion for studies of brain development.

**Figure 2:**
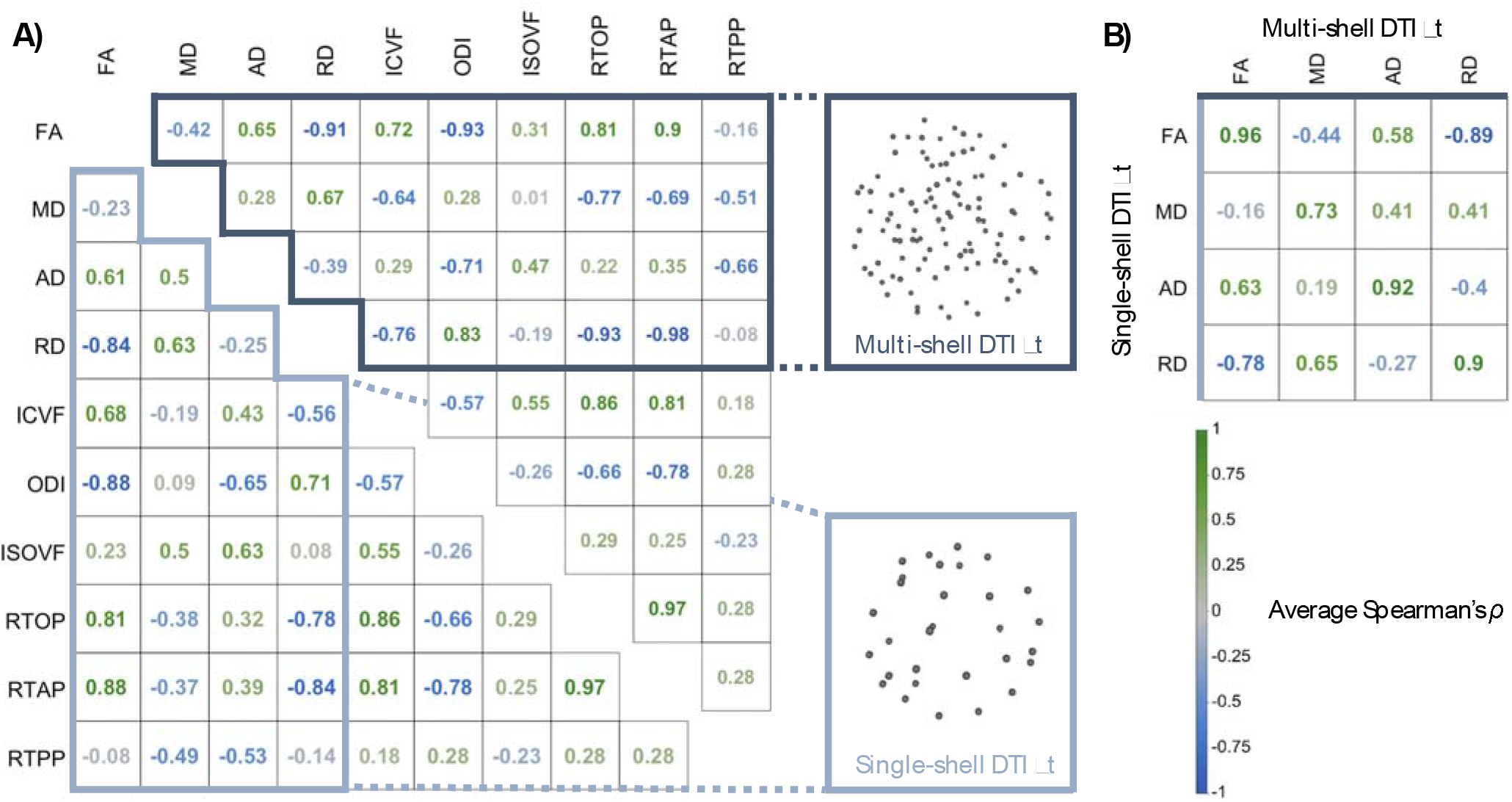
Measures of diffusion are differentially related. **A.** Average Spearman’s correlations between diffusion metrics in white matter. The top triangle depicts correlations derived from multi-shell DTI fitting, and the bottom triangle reflects correlations derived from single-shell fits. **B.** Average correlations between single and multi-shell DTI metrics. FA = fractional anisotropy, MD = mean diffusivity, AD = axial diffusivity, RD = radial diffusivity, ICVF = intracellular volume fraction, ODI = orientation dispersion index, ISOVF = isotropic volume fraction, RTOP = return-to-origin probability, RTAP = return-to-axis probability, RTPP = return-to-plane probability.

### 3.2 Associations with age vary dramatically by diffusion measure

We evaluated the association of each diffusion metric with age. Both mean white matter values and high-resolution mass-univariate analyses at each voxel were conducted. While controlling for sex and in-scanner motion, the mean white matter values of every diffusion metric were related to age in our sample at a nominal (uncorrected) threshold of *p* = 0.05, with the exception of FA calculated from all shells (**Figure 3A, Table 1**). As expected, associations with age were most prominent at the younger end of the age range sampled and diminished in adulthood (**Figure 3B**).

**Figure 3:**
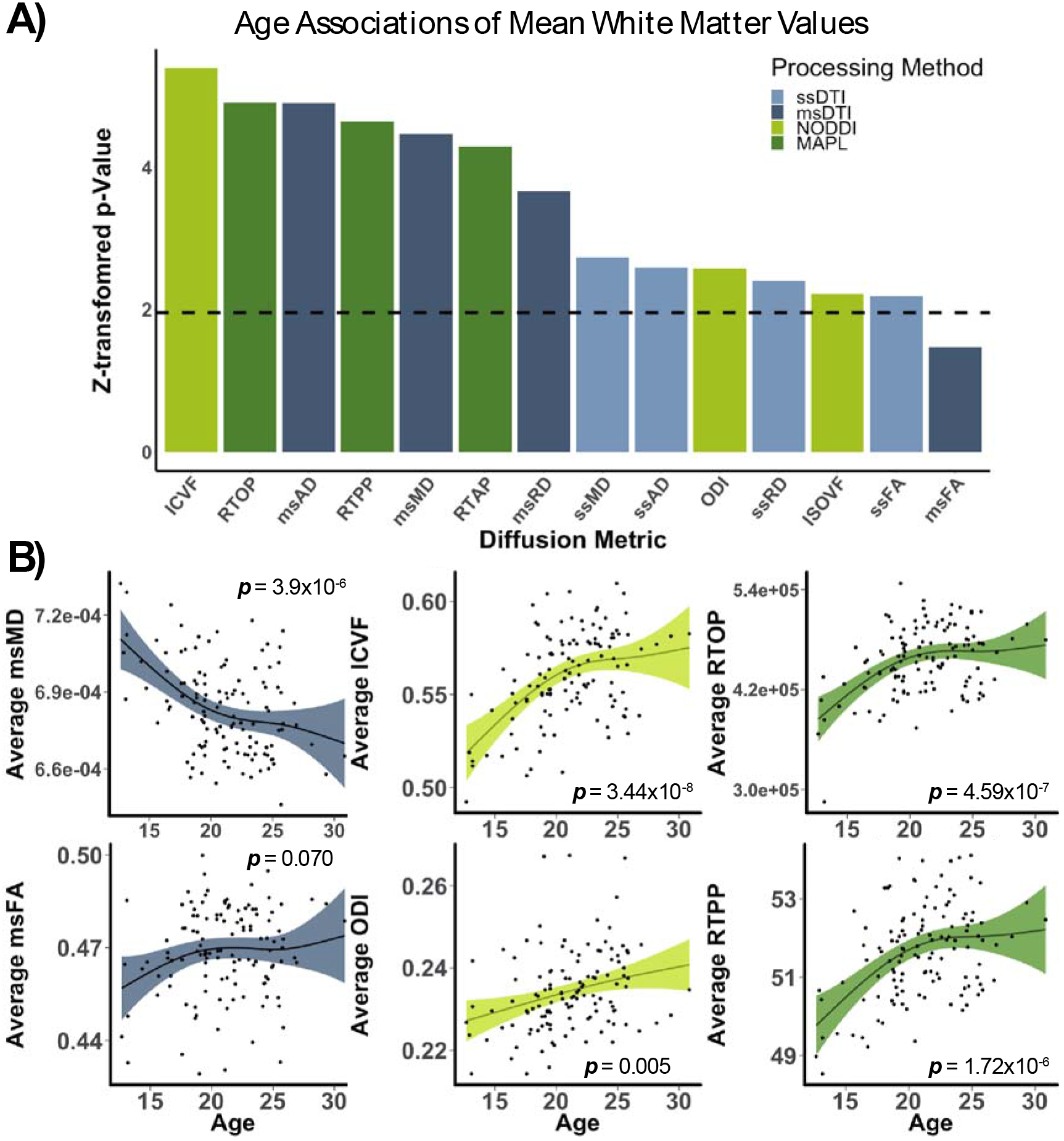
Diffusion models leveraging multi-shell data show variable associations with age in white matter. **A.** Z-scores of *p*-values derived from generalized additive models (GAMs) for each diffusion metric. GAMs were construed with the mean value of each metric as a function of age, with head motion and sex included as linear covariates. The dashed line indicates a nominal, uncorrected significance level (*p* = 0.05, *z* = 1.96). **B.** Relationships between mean white matter values and age, after controlling for sex and data quality. ms = multi-shell, ss = single-shell.

**Table 1:** Mean values in white matter, statistical associations with age, and motion for each diffusion metric. Associations with head motion were evaluated after controlling for age and sex on diffusion metrics.

| Metric | Mean Value | $F_{\text{Age}}$ | $p_{\text{Age}}$ | $r_{\text{relRMS}}$ | $p_{\text{relRMS}}$ |
| --- | --- | --- | --- | --- | --- |
| msFA | .4685 | 2.846 | 0.0702 | -0.241 | 0.0073 |
| msMD | $6.835 \times 10^{-4}$ | 11.68 | $3.98 \times 10^{-6}$ | 0.128 | 0.1589 |
| msAD | $1.073 \times 10^{-3}$ | 14.89 | $4.80 \times 10^{-7}$ | -0.048 | 0.5927 |
| msRD | $4.886 \times 10^{-4}$ | 7.558 | $1.25 \times 10^{-4}$ | 0.191 | 0.0339 |
| ssFA | .4396 | 4.254 | 0.0146 | -.141 | 0.1189 |
| ssMD | $8.219 \times 10^{-4}$ | 5.024 | 0.0031 | 0.201 | 0.0255 |
| ssAD | $1.247 \times 10^{-3}$ | 8.238 | 0.0048 | 0.168 | 0.0626 |
| ssRD | $6.095 \times 10^{-4}$ | 4.911 | 0.0083 | 0.200 | 0.0274 |
| ICVF | 0.5598 | 16.53 | $3.44 \times 10^{-8}$ | 0.098 | 0.2805 |
| ODI | 0.2342 | 5.72 | 0.0050 | 0.343 | 0.0001 |
| ISOVF | 0.0809 | 3.884 | 0.0134 | 0.396 | $5.68 \times 10^{-6}$ |
| RTOP | 454,755 | 13.21 | $4.59 \times 10^{-7}$ | 0.045 | 0.6196 |
| RTAP | 6,493.464 | 10.65 | $8.96 \times 10^{-6}$ | -0.078 | 0.3886 |
| RTPP | 51.6496 | 12.66 | $1.72 \times 10^{-6}$ | 0.0812 | 0.3717 |

Voxelwise analyses within white matter revealed variable patterns. While some metrics demonstrated little only sparse associations with age after correcting for multiple comparisons (FDR *Q*<0.05), RTAP, RTPP, RTOP, ICVF, and MD derived from all of the shells were significantly related to age in several thousand voxels each (**Figure 4**). Sex was not significantly related mean white matter values for any metric evaluated, and less than 80 voxels were significantly related to sex in voxelwise analyses.

**Figure 4:**
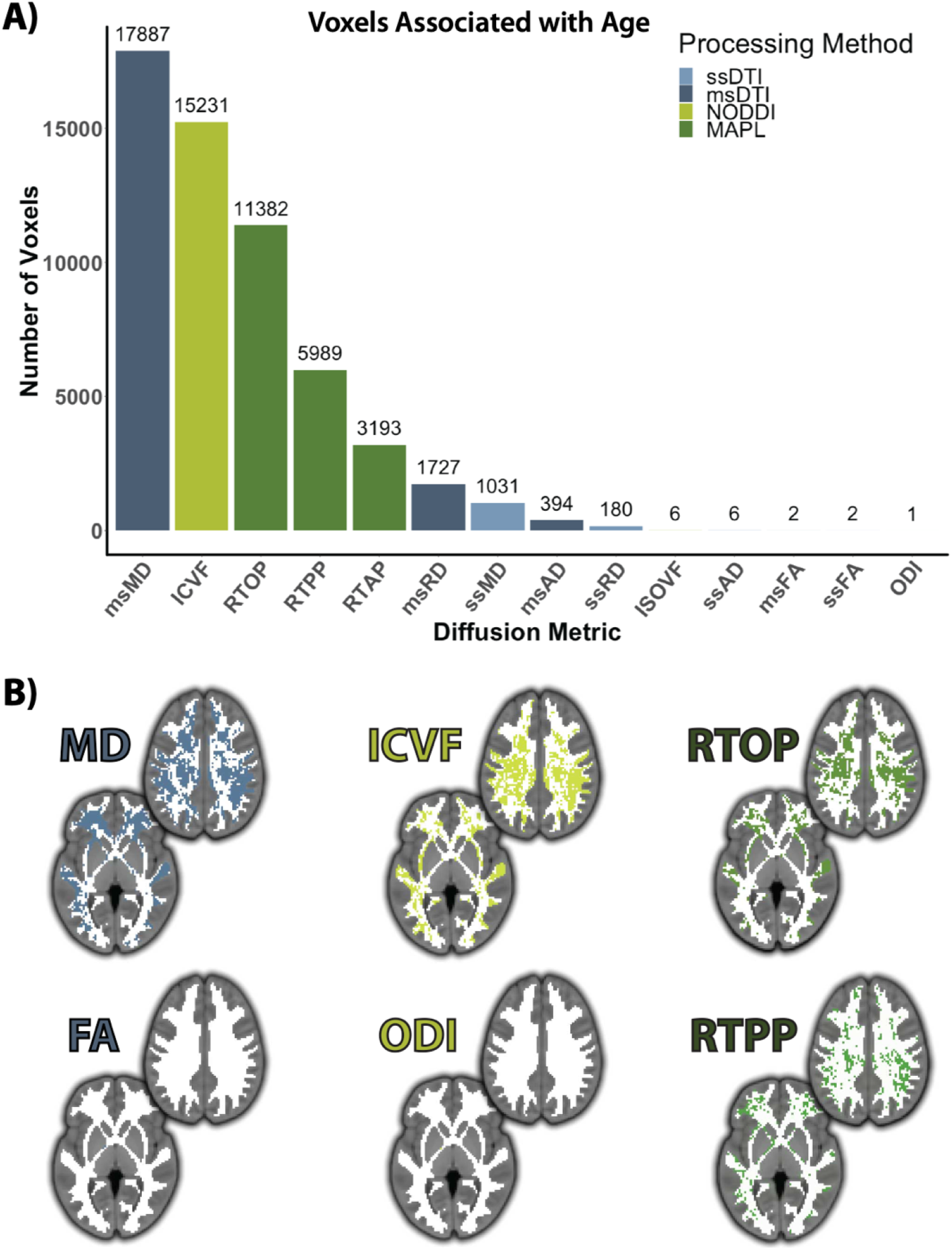
Regional patterns of neurodevelopment are differentially associated with diffusion metrics. **A.** Number of voxels that displayed significant associations with age following mass-univariate analyses while controlling for sex and head motion. Evaluations were corrected for multiple comparisons via FDR-correction (*Q* < 0.05), **B.** Locations of age-associated voxels in the standard space template and white matter mask used for analysis; the mask itself consisted of 48,504 voxels.

### 3.3 Estimates of network development vary according to diffusion metric

Given that tools from network science are increasingly used to study the developing brain, we next evaluated associations with age within networks where edges were weighted by diffusion metrics. These analyses yielded similar results as the voxelwise analyses described above, with network edges weighted by ICVF, MD, RTOP, RTAP, and RTPP displaying the most associations with age (**Figure 5**).

**Figure 5:**
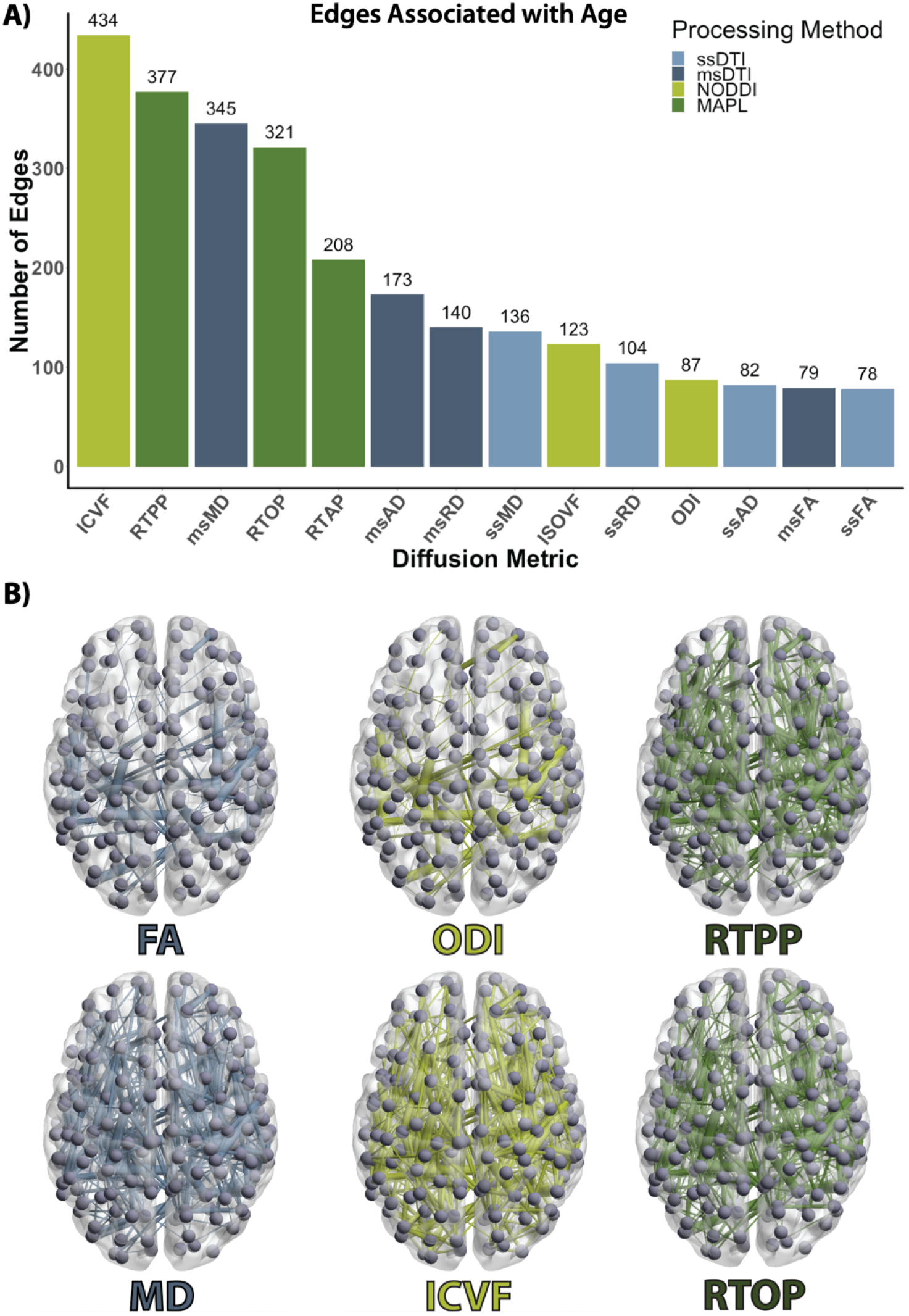
Scalar-weighted structural networks show differential associations with age. **A.** Number of edges that displayed significant associations with age following FDR-correction (*Q* < 0.05), while controlling for sex and head motion. **B.** Associations between age and selected structural networks; thickness of edges is scaled to their *z*-transformed *p*-values, with lower *p*-values depicted by thicker edges, multiple comparisons were accounted for using FDR (*Q* < 0.05).

### 3.4 Diffusion measures are differentially impacted by data quality

As a final step, we sought to characterize the impact of motion on all diffusion metrics. Evaluation of mean white matter values revealed that several mean diffusion measures were significantly related to head motion after controlling for age and sex, including FA, RD, ODI, and ISOVF (**Table 1, Figure 6**). At the voxel level, there were few associations with motion that survived correction for multiple comparisons. After FDR correction, less than 25 voxels were significantly associated with motion for all metrics except ISOVF, which had 239 voxels significantly associated with head motion. Similarly, analyses of networks weighted by each of these values revealed that less than 27 edges were associated with head motion for all three measures, except for ISOVF, which had 77 edges significantly associated with head motion.

**Figure 6:**
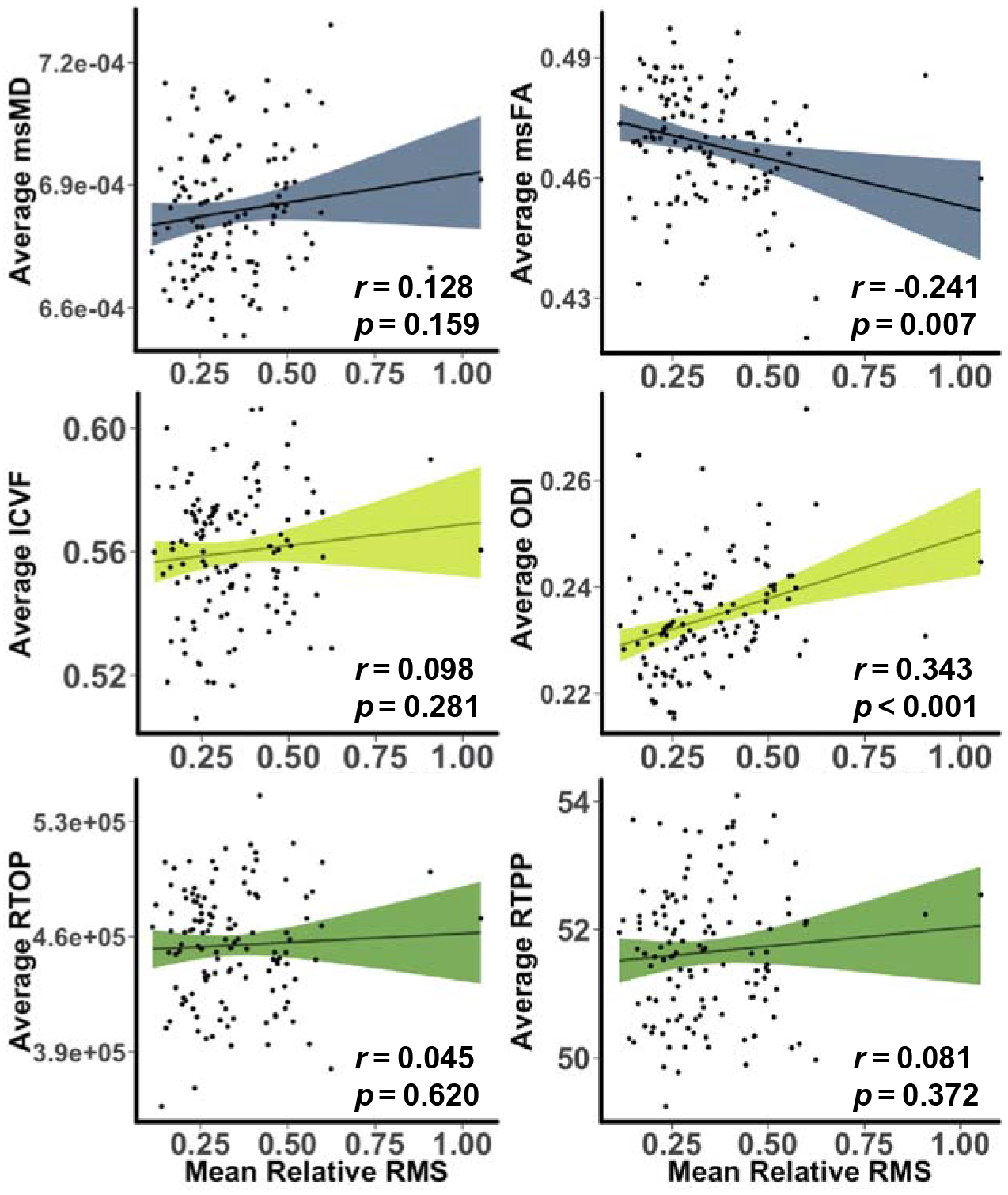
Measures of diffusion are differentially impacted by in-scanner motion. Selected measures displayed; see Table 1 for full results. All analyses control for age and sex.

## 4. DISCUSSION

Our findings suggest that diffusion models leveraging multi-shell data have important advantages for studying the developing brain. These advantages include increased sensitivity to developmental effects and reduced impact of in-scanner motion, which were seen multiple scales, including mean white matter values, voxelwise analyses, and network edges. The context, implications, and limitations of these results are discussed below.

### 4.1 Metrics derived from multi-shell data demonstrate superior sensitivity to brain development

In our dataset, diffusion models that leveraged the full multi-shell acquisition had strong associations with age. Associations with age were seen in measures derived from NODDI (ICVF), MAPL (RTOP, RTAP, RTPP), and multi-shell (AD, MD, and RD). This is particularly interesting for diffusion metrics that were not highly correlated with others, like RTPP, as they may convey unique information regarding neurodevelopmental microstructure changes. In contrast, ODI and ISOVF did not demonstrate substantial associations with age. However, this does not imply underperformance of the NODDI model; there is no strong *a priori* reason to believe that the dispersion of neurite orientations or isotropic water diffusion would dramatically change in white matter over this age range of neurodevelopment.

In contrast to NODDI, MAPL, and DTI fit to all shells, single-shell DTI metrics tended to demonstrate fewer age associations in all analyses. Although these metrics were calculated from less diffusion directions than their multi-shell counterparts, DTI-based neurodevelopmental inquiries have effectively characterized microstructure with far fewer sampling directions (Lebel et al., 2008), indicating that differences between DTI fitting methods may not be driven by the number of directions sampled. However, it is important to consider that the diffusion tensor model does not explicitly account for the non-gaussianity of water diffusion that is common at higher *b*-values.

Despite this shortcoming, most DTI-derived metrics fit using all shells demonstrated substantial associations with age, in particular MD. This indicates that complete fulfillment of the theoretical assumptions underlying the diffusion tensor model may not be necessary for probing broad, albeit potentially unspecific, developmental effects. Finally, the utility of fitting the diffusion tensor model to multi-shell data may have been enhanced in our study due to our acquisition protocol not utilizing *b*-values over 2000 s/mm^2^, which yields data that is most at odds with the assumptions of the diffusion tensor model.

These results move beyond previous findings in several respects. First, to our knowledge, this is the first study to demonstrate that MAPL-derived metrics are highly sensitive to brain development in youth. Second, our results demonstrate that multi-shell measures of local microstructure and structural brain network connectivity such as ICVF and RTPP are more strongly associated with age than traditional FA-weighted networks. This result builds upon prior studies, which have shown that ICVF from NODDI is more strongly associated with age than traditional measures such as FA (Chang et al., 2015; Genc et al., 2017; Ota et al., 2017). Third and perhaps most importantly, these results emphasize the utility of multi-shell modeling techniques for studying brain development. These advantages of multi-shell models likely stem from their ability to successfully capture differential tissue responses across b-values, and the evolution of complex white matter architecture during development (Jeurissen et al., 2013; Volz, Cieslak, and Grafton 2018).

### 4.2 MAPL metrics are less impacted by head motion than NODDI and DTI

As children are more likely to move during scanning adults, motion artifact remains major concern for studies of brain development (Theys, Wouters, & Ghesquière, 2014; Satterthwaite et al., 2013; Fair et al., 2012). For diffusion imaging and other sequences, the primary determinant of scan quality for diffusion imaging is in-scanner head motion (Yendiki et al., 2014; Ling et al., 2012). Importantly, higher in-scanner motion was associated with reduced mean white matter FA, and increased MD, RD, AD, ODI, and ISOVF while accounting for age. This finding aligns with prior reports of in-scanner motion systematically impacting DTI metrics (Yendiki et al., 2014; Ling et al., 2012, Roalf et al., 2016; Baum et al., 2018).

However, to our knowledge there has been no prior work documenting the impact of in-scanner head motion on ODI and ISOVF, or any measure derived from MAPL. ODI and ISOVF were both significantly positively correlated with in-scanner head motion. Investigators should consider and account for this confound when utilizing the NODDI model. Notably, measures derived from MAPL were minimally impacted by motion. This may be due to the Laplacian signal regularization in MAPL, which was designed to mitigate the negative impact of noise in DWI acquisitions. Especially when considered in combination with the robust associations between MAPL-derived measures (RTOP, RTAP, and RTPP) and age, such noise-resistance may strengthen the rationale for using MAPL in studies of brain development.

### 4.3 Limitations and future work

Several limitations should be noted. First, our results were only derived from one scanning protocol and scanner. Replication of these results in other multi-shell protocols would strengthen evidence for the relative advantages of multi-shell models. Second, our study only evaluated adolescents and young adults. Studies of younger children would provide complimentary data, as white matter may undergo even more dramatic microstructural changes at earlier ages, and FA-measured effects would likely be more apparent (Lebel et al., 2008). Indeed, similar inquiries in older populations have unveiled different trends in microstructural changes over aging, indicating that these trends in diffusion metrics may be substantially different in different age ranges (Kodiweera et al., 2016). Third, it should be noted that compared to prior studies (Roalf et al., 2016, Yendiki et al., 2014, Baum et al., 2018), we detected a relative paucity of relationships between diffusion metrics and motion. This may be due to both advances in diffusion image processing techniques (such as the outlier replacement implemented in FSL’s *eddy*) and reduced statistical power. While the sample size of the present study is not small, it is a fraction of the size of other recent reports that evaluated the impact of motion on single-shell acquisitions (Baum et al., 2018, Roalf et al., 2016). Fourth, we used deterministic DTI-based tractography to define the streamlines, which results in a sparse structural network biased towards major white matter tracts. While these network analyses demonstrated enhanced associations with several diffusion metrics, networks constructed using multi-fiber tractography techniques might provide additional advantages (Maier-Hein et al., 2017; Reddy & Rathi, 2016; Farooq et al., 2016; Christiaens et al., 2015; Boniha et al., 2015).

### 4.4 Conclusion

In summary, we provide novel evidence that diffusion metrics are differentially associated with age and motion in youth. Measures that are more tightly linked brain maturation and less related to data quality are likely to be particularly useful for developmental or clinical samples. Through free open-access software, these advanced diffusion methods are relatively easy for investigators to implement (Alimi et al., 2018; Daducci et al., 2015; Fick, Deriche, and Wassermann 2018; Garyfallidis et al., 2014). In the context of these results, we anticipate that multi-shell diffusion models will be increasingly adopted by the developmental and clinical neuroscience community.

## Notes

#### Summary of Updates

We've expanded initial analyses from three metrics of diffusion to fourteen across DTI, NODDI, and MAPL. This includes DTI metrics calculated from the b=800 shell and the entire multi-shell dataset.

